# Interaction of Hippocampal Ripples and Cortical Slow Waves Leads to Coordinated Large-Scale Sleep Rhythm

**DOI:** 10.1101/568881

**Authors:** Pavel Sanda, Paola Malerba, Xi Jiang, Giri P. Krishnan, Sydney Cash, Eric Halgren, Maxim Bazhenov

## Abstract

The dialogue between cortex and hippocampus is known to be crucial for sleep dependent consolidation of long lasting memories. During slow wave sleep memory replay depends on slow oscillation (SO) and spindles in the (neo)cortex and sharp wave-ripple complexes (SWR) in the hippocampus, however, the mechanisms underlying interaction of these rhythms are poorly understood. Here, we examined the interaction between cortical SOs and hippocampal SWRs in a computational model of the hippocampo-cortico-thalamic network and compared the results with human intracranial recordings during sleep. We observed that ripple occurrence peaked following the onset of SO (Down-to-Up-state transition) and that cortical input to hippocampus was crucial to maintain this relationship. Ripples influenced the spatiotemporal structure of cortical SO and duration of the Up/Down-states. In particular, ripples were capable of synchronizing Up-to-Down state transition events across the cortical network. Slow waves had a tendency to initiate at cortical locations receiving hippocampal ripples, and these “initiators” were able to influence sequential reactivation within cortical Up states. We concluded that during slow wave sleep, hippocampus and neocortex maintain a complex interaction, where SOs bias the onset of ripples, while ripples influence the spatiotemporal pattern of SOs.

## Introduction

Coordination between thalamo-cortical and hippocampal (TH-CX-HC) networks during slow-wave sleep is implicated in the process of memory consolidation. The theory of two stage memory formation [Squire and Alvarez, 1995] assumes that newly acquired memory traces created during recent experience initially depend on the hippocampal structures but become hippocampus independent during the following stage of consolidation [McClelland et al., 1995, Frankland and Bontempi, 2005]. Hippocampus may still preserve an index code to link together elements of more complex memories [Teyler and DiScenna, 1986, Nadel et al., 2007, Winocur et al., 2010]. The underlying mechanisms mediating memory consolidation during sleep are not well understood, but hippocampal sharp-wave ripples (SWR) coordinated by the cortical slow oscillations (SO) were shown to participate in the consolidation process [Girardeau et al., 2009, Nakashiba et al., 2009, Ego-Stengel and Wilson, 2010, Wang et al., 2015]. Indeed, a complex nesting of different sleep graphoelemensts was recently reported in vivo [Staresina et al., 2015, Latchoumane et al., 2017]. While a phase preference for SWR with respect to ongoing SO was reported in several studies [Sirota et al., 2003, Isomura et al., 2006, Mölle et al., 2006, Peyrache et al., 2011], SWR complexes can be detected at any SO phase and it remains unclear if SWRs happening at different phase of the SO cycle are performing different functions [Maingret et al., 2016]. In this new study, we ask two related questions: how ongoing SOs affect ripple occurrences and vice-versa how ripples shape the spatiotemporal patterns of Up and Down cortical states, the alternating activity in cortical neurons underlying sleep SO [Steriade et al., 1993b, Sanchez-Vives and McCormick, 2000].

To study the interaction of SWRs and SOs, we bring together biophysical models of the thalamo-cortical network [Bazhenov et al., 2002, Krishnan et al., 2016, Wei et al., 2018] which reproduces SO-like activity during NREM stage-3 sleep, and hippocampal CA3-CA1 circuitry producing sharp-ware ripple events [Malerba et al., 2016, Malerba and Bazhenov, 2018]. Both networks were connected within a cortico-hippocampal synaptic feedback loop. We observed that the cortical input was driving SO-ripple coupling. At the same time, ripples influenced the structure of SOs subtly – depending on the phase of SO, ripple could either anticipate or postpone transitions between Up and Down-states, as well as change the initiation site and synchronization properties of the slow waves in the population of cortical neurons. We also observed that a cortical site receiving ripple input at a given cycle of SO, would influence the cortical spatiotemporal pattern in subsequent SO cycles. Finally, we show that the SO-ripple interaction can influence cortical synaptic plasticity, and hence shape sequential spike reactivation among cortical cells, as reported previously *in vivo* [Euston et al., 2007, Ji and Wilson, 2007, Peyrache et al., 2009].

## Results

### Organization of the network

SO are generated in the thalamocortical network [Steriade et al., 1993b,a, Timofeev et al., 2000, Volgushev et al., 2006, Mohajerani et al., 2010, Sheroziya and Timofeev, 2014] and significantly interact and coordinate with hippocampal SWRs [Buzsáki, 2015]. Our model builds up on the two major network blocks (see Fig. 1) – an oscillating thalamocortical (TH-CX) network generating slow (~0.7 Hz) cortical oscillations [Bazhenov et al., 2002, Krishnan et al., 2016, Wei et al., 2018], and a hippocampal (HC) network spontaneously generating SWRs with an average oscillation frequency within ripples ~155 Hz [Malerba and Bazhenov, 2018]. The single layer of cortical neurons (CX) displays alternating Up and Down states and is further synchronized by the activity of thalamic TC-RE cells [Lemieux et al., 2014]. The connectivity between cortex and hippocampus in the model resembles the biological circuitry where global Up-states tend to travel from medial prefrontal cortex to the medial temporal lobe and hippocampus [Nir et al., 2011]. The output side of hippocampal processing – CA1/subiculum - then projects one of its streams back to mPFC via the fornix system [Cenquizca and Swanson, 2007] (apart from major feedback connectivity back to the entorhinal cortex, which we do not model here). Thus, in our implementation a restricted region of the cortical network projects to a subset of CA3 cells in hippocampus and affects the probability of sharp-wave generation there. CA3 region is consequently connected to the CA1 region which displays ripple events as a consequence of large excitatory events occurring in CA3. The major part of CA1 output is then fed back to the cortical region opposite to the region projecting back to CA3. For specific details of the connectivity see Methods. We did not specifically tune the network so that the Up-state would start at a specific region, however, as we show later, the ‘frontal’ part of the cortex tends to start Up-states as a result of CA1 output activity targeting this region.

**Figure 1:**
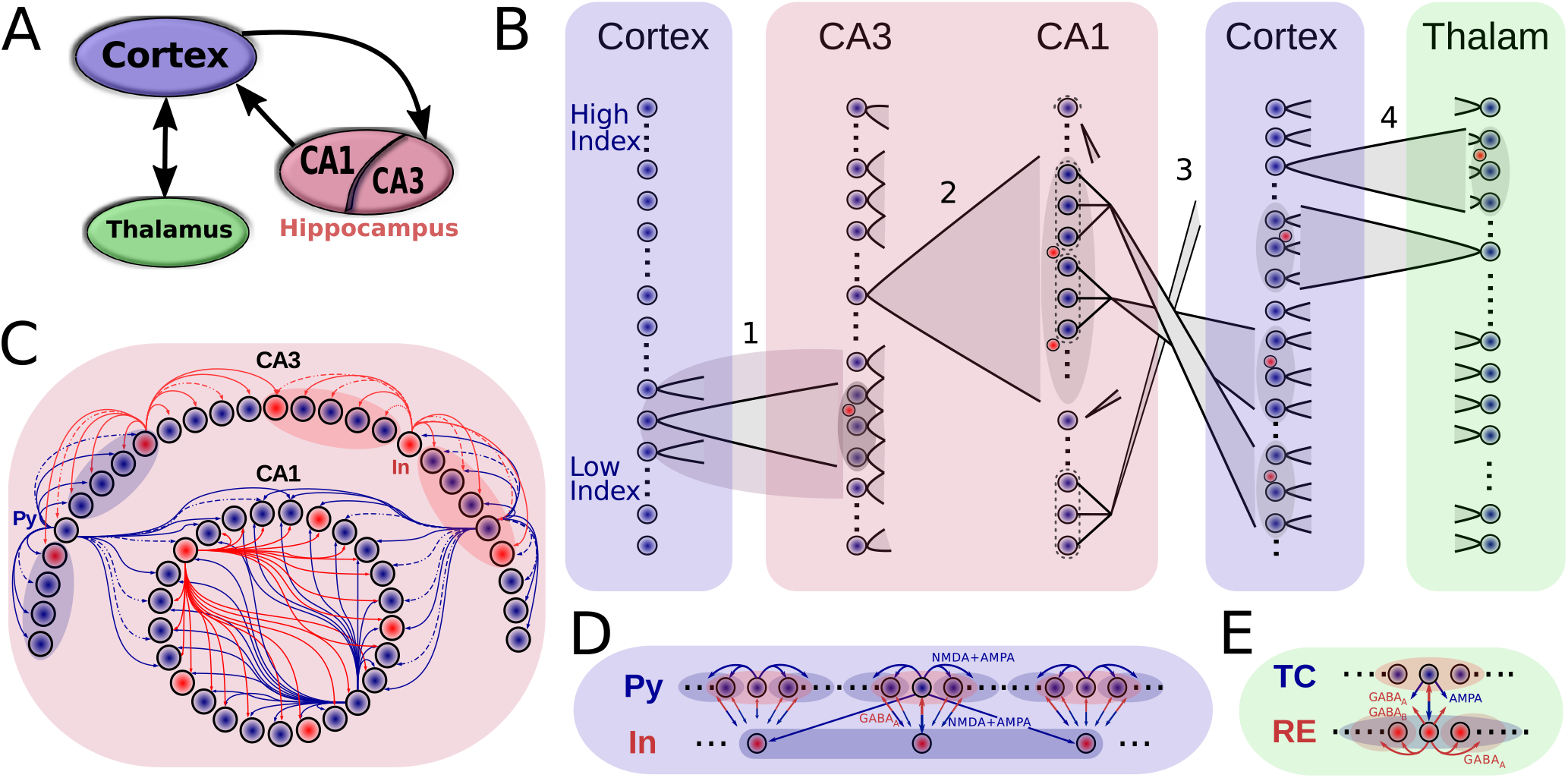
Model connectivity. A. The model consists of the thalamocortical loop which generates slow oscillations (SO), and the hippocampal circuit (consisting of CA1 and CA3 regions) which generates sharp wave - ripples (SWR). The two components are connected via cortical input to CA3 and hippocampal output from CA1. B. Details of network connectivity. (1) Cortex->CA3: a small contiguous population of cortical excitatory cells targets a restricted part of CA3 which is highly responsive to incoming excitation. Both CA3 excitatory (blue dots) and inhibitory cells (red dots) were targeted. (2) CA3->CA1 (Schaffer collaterals). CA3 pyramidal cells broadly target CA1 E/I cells. (3) CA1->Cortex. Each small patch of CA1 cells project to a small focal region in Cortex. (4) Cortex<->Thalamus. Cortical pyramidal neurons target both thalamic RE and TC cells, TC cells project back to both pyramidal and inhibitory interneurons of cortex. Cells in each region are linearly arrayed, with connectivity between regions generally between cells at the same level, with the sole exception of the CA1-> Cortex where HC cells at the top of the array project to cortical cells a the bottom, and vice versa. C-E. Intra-area connectivities. Blue circles/arrows are excitatory cells/connections, the red are inhibitory ones. The shaded area designates the target region of a projecting neuron. Connectivity of the thalamocortical circuity closely follows [Wei et al., 2018], hippocampal connectivity is similar to the connectivity used in [Malerba and Bazhenov, 2018].

Fig. 2 shows the spiking rastergrams and LFPs of all the network components when coupled into a “close-loop” large-scale network. The most prominent hippocampal features are sharp-wave events internally generated in CA3 and ripples in CA1 (LFP is filtered in the high frequency range for better visibility of the ripples). By design, the neurons in the bottom region (i.e. with low index number) of CA3 had higher strength of lateral connections and thus were more prone to be part of a sharp-wave event (note Fig. 2, top right panel, with majority of excitatory spikes located in the bottom region during the sharp-wave). Due to the connectivity from CA3 to CA1, CA3 SWs triggered CA1 ripples in the topologically equivalent (in terms of cell indexes, see Fig. 1B) region of the CA1 (Fig. 2, second right panel). Fig. 2, left top, shows detailed view of a single ripple event and a histogram of probability of CA1 cell to become part of a ripple. In order to avoid a problem that the same region of cortex both receives the majority of ripple events from CA1 (see histogram in Fig. 2 top left) and targets CA3 network, CA1->CX connections were initially “flipped” (see Fig. 1B), such that the low CA1 region projected to the upper region of the cortical network and vice versa (we consider network model without flipping in later sections).

**Figure 2:**
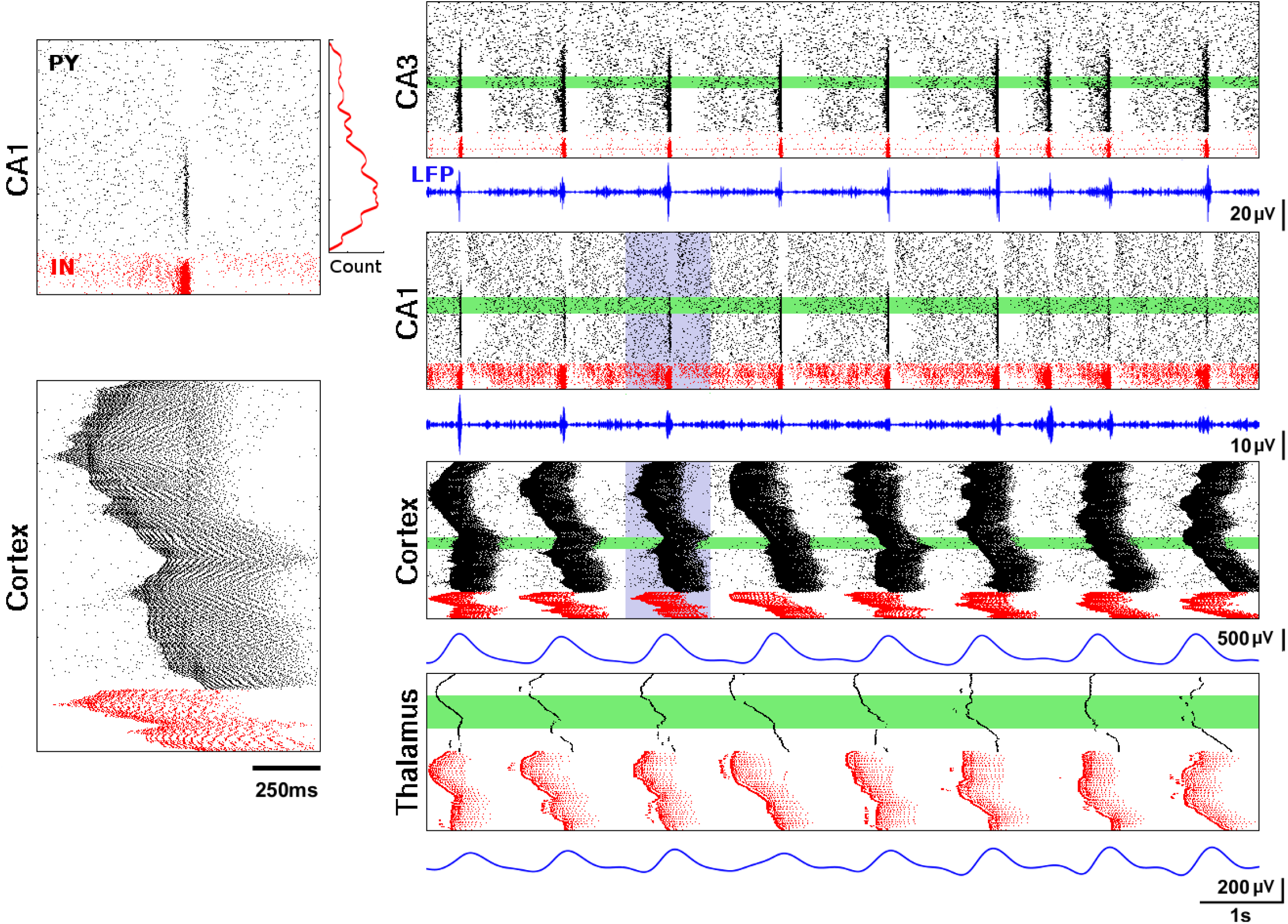
Close-loop network dynamics. Spiking activity of pyramidal (black) and local inhibitory cells (red), LFP traces (blue). Left panels: representative ripple event in CA1 (top) and Up-state in cortical network (bottom), correspond to the violet region on the right. Histogram next to the ripple event shows average spike count of CA1 excitatory cells during ripples. Right panels from top to bottom: 10 seconds of full network activity; spiking rastergrams for CA3, CA1, cortical network, and thalamic network (RE and TC cells); average LFPs from 100 neurons (green areas). LFP for CA1 was filtered from 120 to 200 Hz, for CA3 from 90 to 200 Hz, and for both cortical and thalamic cells from 0.5 to 2 Hz.

The cortical network generated a regular slow oscillatory pattern (Fig. 2, third right panel) in which pyramidal cells alternated between Up and Down-states with an approximate frequency of 0.7 Hz. A single Up-state is detailed in Fig. 2, left bottom, and shows nested oscillatory spindle-like activity sequentially traveling throughout the network.

### Coordination of SO-SWR in model and experimental data

To measure coordination between the network activities we used three reference points – Up-to-Down transition (UDt), Down-to-Up transition (DUt) and SWR event time – and we compared model simulations with experimental data. Recordings were done in long-standing drug-resistant epileptic patients with intracranially implanted electrodes (see details in Methods). Two representative electrodes were chosen - one in the hippocampus (Fig. 3A, left) and one in prefrontal cortex (Fig. 3A, right). Recordings from the NREM phase of sleep were analyzed. SO were detected in the frontal electrode (example shown in Fig. 3B, top), while SWR were detected in the hippocampal electrode, see example in Fig. 3B, bottom. We compared the locking between SWR and transition points of slow rhythm for both model and experimental data. Several features were observed. First, there was increase of transitions from Up to Down state after SWR (Fig. 3C, top), but not from Down to Up (Fig. 3C, bottom). Second, number of SWRs increased after transition to Up state (Fig. 3D, bottom panels), but not after transition to Down state (top panels). The peak of SWRs in 3C, top corresponds to peak of UDt in 3D, top. These empirical results were well captured by the model (compare Data and Model panels in each plot). However, we observed more significant variations in number of events in the model vs recorded data. This can be explained by lower baseline activity in model simulations and it can be a results of much more synchronized activity between directly connected cortical and hippocampal areas in the model.

**Figure 3:**
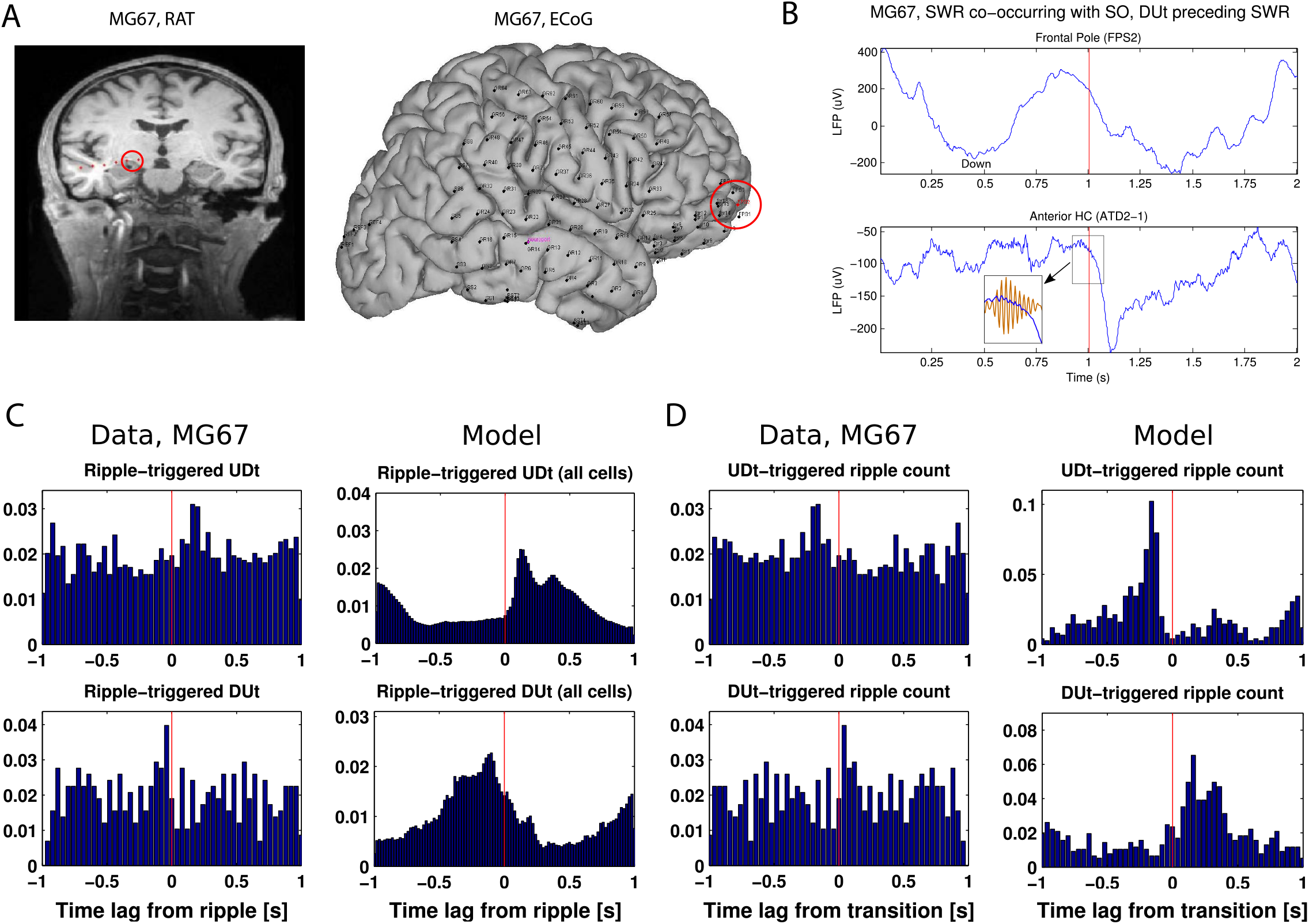
A. Position of electrodes on the subject MG67. Left. Hippocampal depth electrodes image (RAT). Right. ECoG electrodes positions on the same subject. B. Top. Raw LFP trace from ECoG, frontal pole electrode (FPS2, red circle at panel A, right) showing alternating Up/Down states. Bottom. Raw LFP trace from depth electrode (in bipolar montage) from the vertical part of the hippocampal head (red circle on panel A, right) showing ripple following DUt at the top panel. The inset shows signal filtered at the range of 80-120 Hz showing the ripple. Both panels are centered at the ripple (red vertical line, highest point in the analytical amplitude envelope of the filtered signal). C&D. Comparison of model (20 independent trials, each simulating 50s of real time) and experimental data (~16 h of NREM sleep). C. Ripple-triggered DUt/UDt histogram for the model and in the signal from electrodes selected in the panel A. To get a smoother distribution, the transition event in each cell is measured separately in the model; global transition in LFP is used for experimental data (see methods) D. DUt/UDt-triggered ripple histogram for the model and in the signal. Global transition is used for both model and data. Red vertical line aligns with *t* = 0 ms.

### Global coordination of SO-SWR rhythms is set by cortical drive

Next we investigated the mechanisms driving the relationships between cortical and hippocampal events reported in the previous section. In Fig. 4A1 we again plot the DUt-triggered ripple-count histogram in which most of the ripples follow onset of an Up-state. To find what determines this relationship, we considered two open-loop model configurations – one where only CX targets HC and no connections are fed back from HC to CX (column B) and another one where only HC targets CX (column C). As Fig. 4B1 shows, the network model with CX targeting HC preserves the phenomenon of DUt preceding the ripple event. In contrast, in the model where HC targets CX but no feedback projections were implemented, no obvious coordination pattern was observed (Fig. 4C1). We thus conclude that the global coordination of rhythms is set by the cortical drive to the HC network. Nonetheless, some differences to the closed loop model were observed. Removing HC->CX projections simplified the distribution that only revealed one peak (compare Fig. 4 B1 and A1). Two sample Kolmogorov-Smirnov test comparing the distributions of time-lags confirmed that the lags were derived from different distributions, p<0.001, suggesting different behavior of the closed loop model especially near the time of UDt, presumably caused by CA1 activity. Analysis of the ripple event histogram using UDt as a reference point found a sharp peak of ripple activity immediately preceding UDt (Fig. 4A2), while this peak was significantly advanced and less visible in the model missing CA1 input (Fig. 4B2, a Kolmogorov-Smirnov test confirmed that the lags were derived from different distributions, p<0.001).

**Figure 4:**
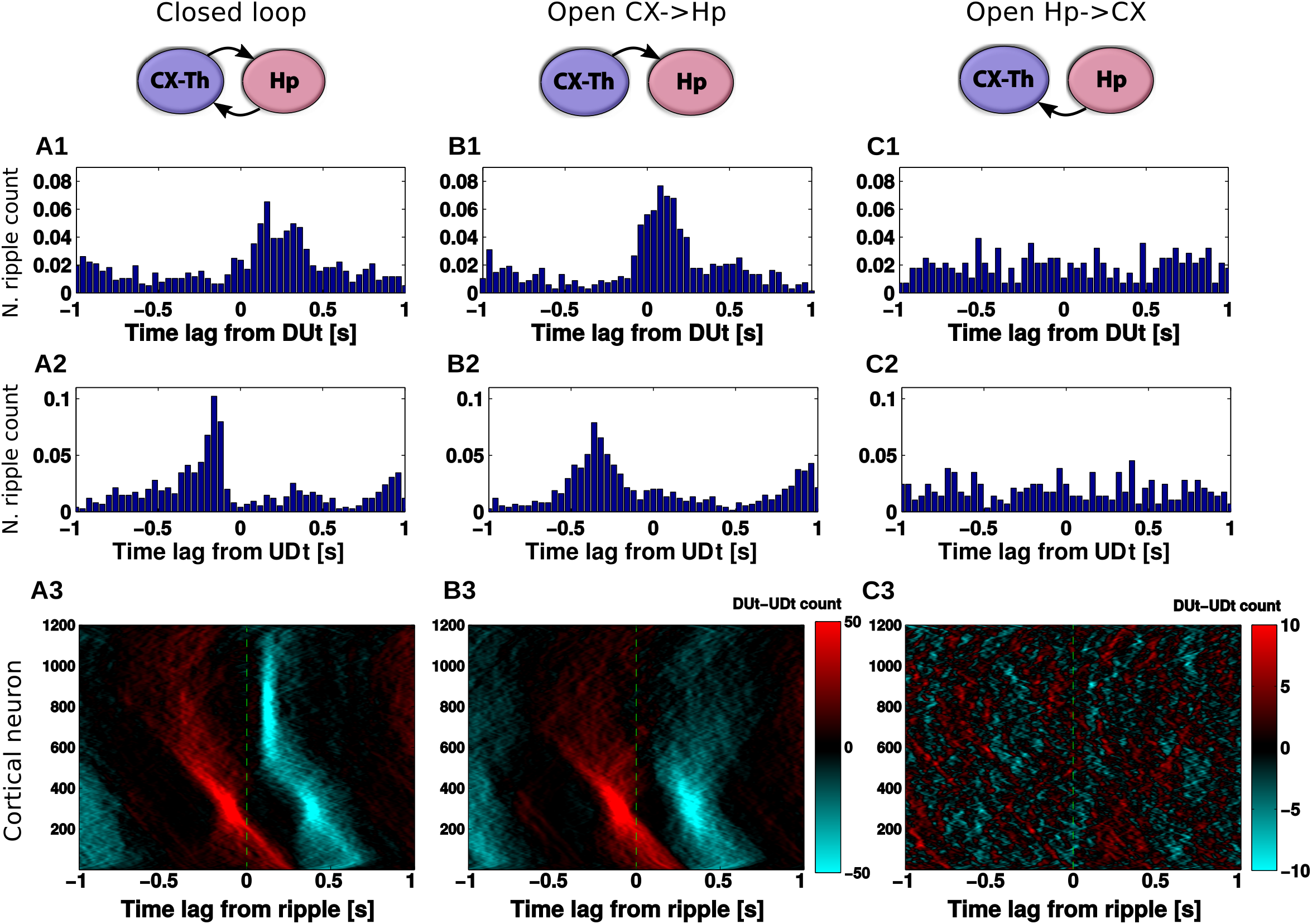
Effect of network structure on DUt/UDt-ripple coordination. Left column (A): closed CX-HC loop, middle column (B): open loop (HC does not project back to CX), right column (C): open loop (CX does not project back to HC). 1: DUt triggered ripple count. 2: UDt triggered ripple count. 3: Spatiotemporal profile of ripple triggered DUt (red) and UDt (blue) count for each CX cell separately. Colormap: DUt-UDt count (red indicates mainly DUt events, blue – mainly UDt events); y-axis - index of CX cells. Note the different scale used for C3. Corresponding cumulative histograms are shown in SI Fig. 7. All averages are across 20 trials.

To further explore the interaction between SWR and Up/Down state transitions, we plotted ripple-triggered DUt-UDt count histograms for each cortical cell separately (Fig. 4A3, B3, C3, the cumulative histograms are shown in SI Fig. 7). Both Fig. 4 A3, B3 show similar UDt/DUt activity in the bottom region of the network (low indexes). This highly consistent DUt activity (red dots – high count of DUt events) can be explained by the fact that a DUt transition increases the probability of the ripple events, and CA3 receives input from CX in this region (Fig. 1B, left). Therefore, ripples could likely be triggered by DUt events. Consistent UDt activity (blue dots – high count of UDt events) in the same network region would then follow from the fact that Up-state duration tends to be consistent across different cycles of SO. What makes Figs. 4 A3, B3 visibly distinct is the pattern of events in the top region (high cell indexes). This pattern is very consistent in the closed-loop network (Fig. 4A3), where UDt (blue dots) typically followed a ripple in the region receiving most of the CA1 input. This structure disappeared in the open-loop network (Fig. 4B3) suggesting that hippocampal input to CX was able to synchronize the UDt events across neurons and across many cycles of SO. In the following section we will discuss the mechanism that may lead to increased synchrony of UDt in the closed loop network model.

### Hippocampal influence on the slow oscillation

To better understand the impact of a ripple event on the dynamics of the cortical slow-wave we tested a simplified scenario. The thalamocortical network was simulated in isolation with an input that was identical to a single hippocampal ripple (the exact spiking of CA1 neurons was saved from simulation of the full TH-CX-HC model) and that was applied at different times corresponding to different phases of SO. Thus, for each trial, the cortical network was targeted only at one specific phase of slow oscillation. We ran a set of such trials that uniformly spaced a full cycle of SO and we repeated this for many cycles of SO to get the average response. Fig. 5A shows an example of six trials when stimulation was applied at a different phase to the same cycle of SO (so as to allow for a direct comparison between trials). Cortical spiking before (or after) ripple injection is labeled in black (or blue). The red spikes interposed between blue and black spikes show the directly driven cortical firing in the region receiving HC input. Comprehensive animation of this experiment is provided in SI Fig. 8. As we performed this experiment for many cycles of SO, several consistently repeating phenomena could be observed. First, the ripple arriving at the very end of the Up-state was capable of increasing its duration (compare Fig. 5 A5 vs A1). Second, the ripple arriving approximately in the middle of the Down-state phase was capable of shortening that Down-state and to cause DUt to start sooner (compare Fig. 5 A3 vs A1). Finally, the ripple arriving in the middle of the Up-state phase was capable of improving synchronization of the DUt and mildly shortening it (compare Fig. 5 A6 vs A1). We confirmed these observations by quantification across many different simulations of cortical dynamics (Fig. 5 B-D), including (1) an increase of Up-state duration (the peak in Fig. 5B), (2) a decrease of Down-state duration (the dip in Fig. 5C) and (3) a decrease in the standard deviation for UDt times across neurons in the region receiving the ripple input (the dip in Fig. 5D).

**Figure 5:**
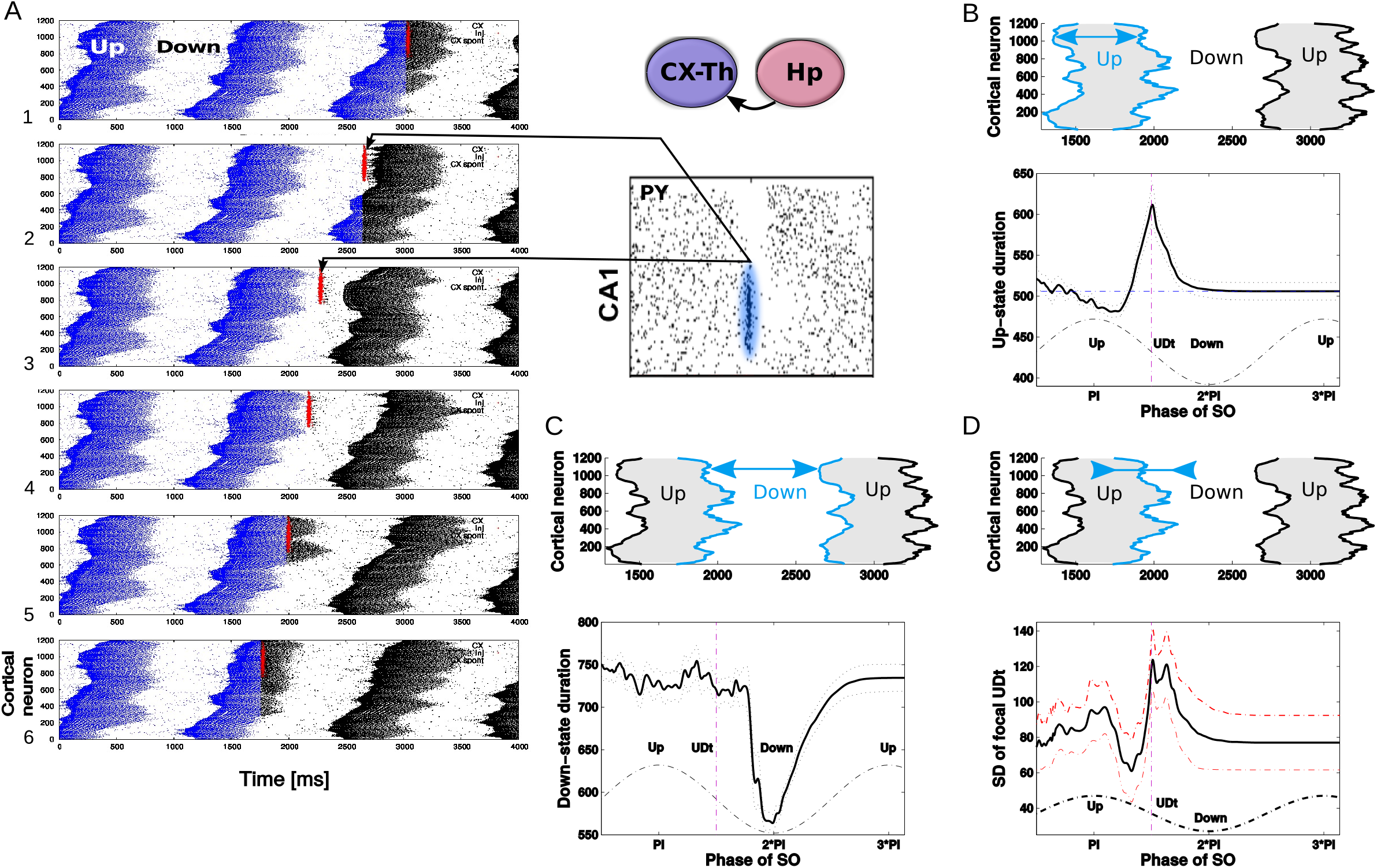
Ripple effect within a single SO oscillatory cycle, open loop scenario. A. A trace of a single representative ripple event was saved from the closed loop simulation and delivered to the isolated thalamocortical network at the different phases of the SO oscillatory cycle. Red dots show spikes of CA1 cells projected to the region of the cortical network. Blue dots are spikes of cortical cells in the network without ripple stimulation, black dots are cortical spikes after the ripple was delivered. The effect of a ripple on the spatiotemporal pattern of DUt/UDt transitions depended on the exact timing of the event. We tested 100 independent trials with identical networks (each trial started with the same initial values of all variables) with the ripple delivered at different times: *T_i_* = *i* * 20 ms for i-th trial. Each trial was repeated 10 times using different initial seed values hereby creating different cortical dynamics; the results were averaged. Animation showing the described experiment is shown in SI Fig. 8. To focus on the effect of a ripple, duration/synchronization effects and SO oscillatory cycle were defined by cortical activity in the region receiving most of the ripple input (top 601-1200 cortical cells). B. Effect of ripple on the Up-state duration. Top. Schematic diagram of cortical activity showing two Up-states (shaded) and single Down-state. Duration of the first Up-state (blue envelope lines) was measured for each trial (different stimulation phase). Bottom. Average effect on Up-state length from 10 runs. X-axis - timing of ripple rescaled to SO cycle (reference cycle for each trial was defined by the run where no ripple was delivered). C. Effect of ripple on the Down-state duration. Top. Schematic diagram of of cortical activity. Duration of the Down-state (blue-line envelope) was measured. Bottom. Average effect on the Down-state duration from 10 runs. D. Effect of ripple on the synchrony of the UDt events. Top: Schematic diagram of cortical activity. Timing of the UDt events (blue-line envelope) across population of cortical neurons was measured. Bottom: Average effect of ripple on the synchronization of Up-state termination measured as a standard deviation of UDt events timing in the cortical neurons.

The dip in the midst of a Down-state (Fig. 5C) suggests that ripples arriving during the middle or later phases of a Down-state (note that Down-state duration curve is skewed to the right) can increase the probability of a network transition to an Up state. This effect was not visible in the full closed-loop network analysis (Figs. 3,4), likely because of a sharp increase in the ripple probability during Up-states tends to hide the relatively small number of ripples that occurred during Down-states. In other words, since most of the ripples occurred during the Up-state, statistically a single ripple was much more likely to be followed by a Down-state than an Up-state; note however a small peak in DUt probability ~200 msec after the ripple in Fig. 3C.

The dip in the midst of Up-state and the peak around UDt in Fig. 5D correspond to the dip/peak in Fig. 5B, suggesting that they can also be interpreted as the ripple occurring during an Up-state generally promoting a synchronous transition to the Down state except when precisely targeted at the very end of a Down state when it can extend its duration (see animations in SI Fig. 8). The increase in synchrony of UDt across cortical neurons during this simplified open-loop experiment suggests the possible cause of the network synchronization observed in the close-loop model. For example, the highlighted blue region in Fig. 4 A3 shows synchronized UDt in the area of the network receiving ripple input (~#600-1000) following a ripple triggered in the late part of Up-state.

The last observation from simplified model was, furthermore, confirmed by running the following experiment. In complete closed-loop TH-CX-HC simulation we varied the delay from CA1 to CX pyramidal cell, thus effectively changing the phase of SO at which cortex tends to receive a ripple. As a result, we observed that the pronounced (synchronized) structure of UDt in the top region of the network (blue dots in Fig. 4A3) appeared mainly for short synaptic delays between CA1 to cortical pyramidal cells (see SI Fig. 4 second/third column, row 1) but disappeared for longer delays (SI Fig. 9 second/third column, e.g., row 3). It is worth noting that UDt-triggered ripple-count histogram in the scenario of shorter delays reveals a peak of ripples ahead of UDt (see Fig. 4A2) suggesting “ripple as a cause of the Down-state”. However, at least in the case of our model, it was mainly the synchronization of transition times to the Down state across population of cortical cell rather than transition to the Down-state itself (which would occur nevertheless just with higher dispersion) which created the sharper peak in the ripple distribution.

The observation that ripples arriving in the midst of a Down-state shortened its duration (“triggered DUt”) from the simplified model (Fig. 5) was also found in the closed-loop model (SI Fig. 4). When CA1->CX axonal delay became (unrealistically) long so that most of the ripples tended to arrive at the midst of the Down-state following the Up-state (which helped to initiate the sharp-wave events), we observed a highly synchronized region of DUt (SI Fig. 4, first column, e.g., 4th row, red dots) initiated by the ripple event occurring in the late phase of the Down-state.

### Ripples influence initiation probability

We describe above how ripple events targeting cortical neurons during specific phases of the slow oscillation are capable of influencing the structure and duration of Up and Down-states. To better understand the cumulative effect of the ripples targeting a specific region of the cortical network, we considered two versions of synaptic wiring differing in the connectivity pattern between the hippocampus and cortex (Fig. 6 A1, B1). This included the original wiring (Fig. 6A1) in which a majority of the hippocampus-triggered ripple output targets the region of cortex distant from the cortical region projecting to CA3 (and thus influencing sharp-wave generation) – mirror mapping - and contrasted it with non-mirror (i.e. direct) mapping, in which the ripple-active CA1 region projects to roughly the same region which targets CA3 (Fig. 6 B1). Simulations of both models revealed that the region receiving most of the ripple input tended to become (after sufficient time; 50s of simulated time in total) a global initiator of the cortical Up-states. Figures 6 A2/B2 show histogram profiles of the global Up-state initiation (two sample Kolmogorov-Smirnov test comparing the distributions of global initiators confirmed that they were derived from different distributions, p<0.001). The inset heatmap in Fig. 6A2 shows that this bias in the initiation preference (yellow region) rapidly decreases as the connectivity strength of CA1->cortical connections (X-axis) decreases. The mean spatial profile of the DUt is displayed as a heatmap in Fig. 6 A3/B3 where for each DUt “wave” a zero time lag indicates that it is the global ignition site (i.e. the first cortical cell which passes through the DUt transition in a given SO cycle). Since in the mirror mapping model Up-states are more likely to start in the top region of the cortex (and conversely in the direct-mapping model they tend to start in the bottom region), this defines a preferred direction of the traveling DUt wave. This observation was further confirmed by measuring an average gradient of the traveling wave for each local neighborhood of neurons that revealed an opposite traveling wave direction for the two network topologies (Fig. 6 A3/B3, histograms). This phenomenon was robust with respect to the phase of SO being hit by the ripple, see SI Fig. 9, 4th column, since varying the time delay of CA1->CX projections did not change the overall shape of DUt traveling wave.

**Figure 6:**
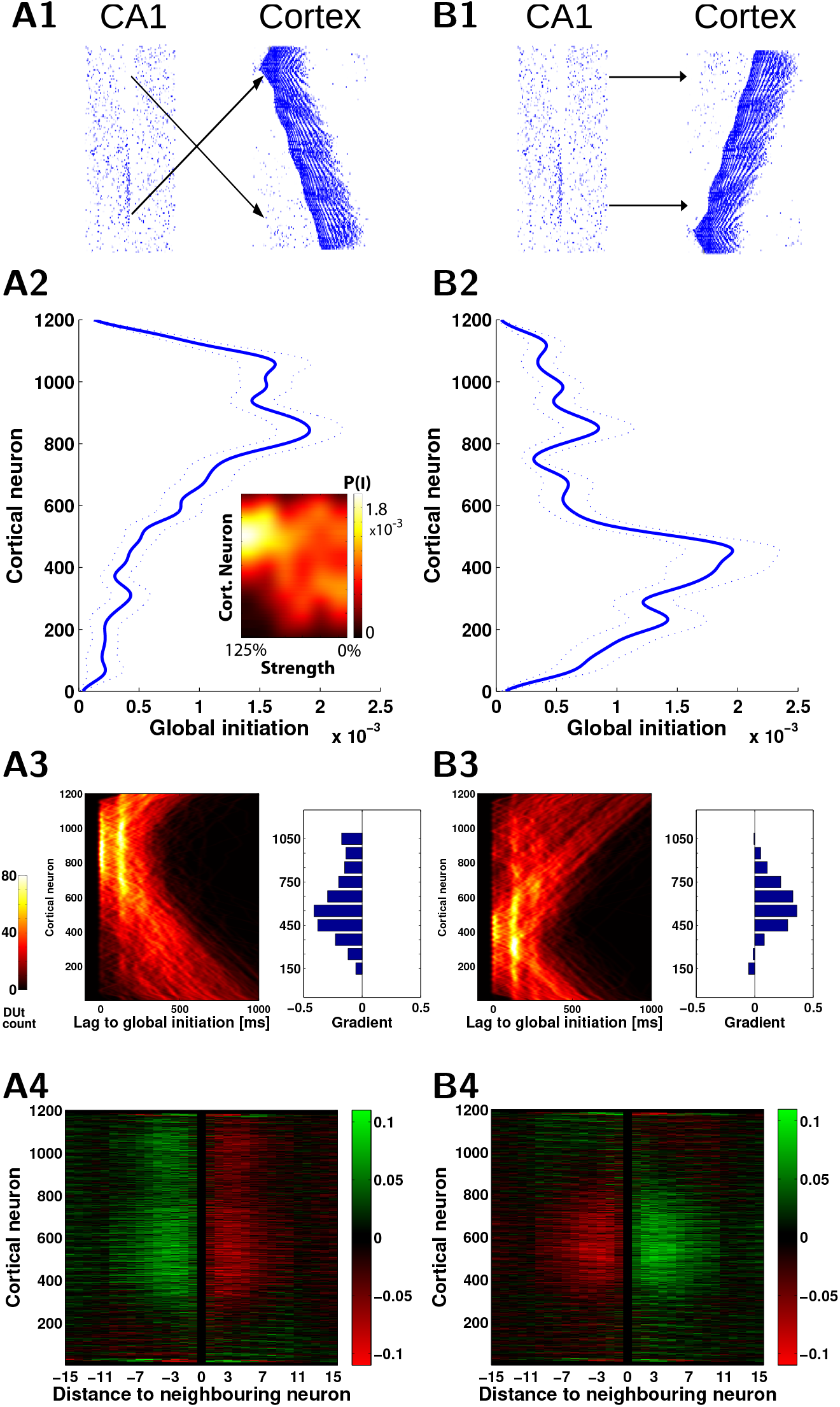
Hippocampal ripples shape slow wave spatiotemporal pattern. A1, B1. Two wiring models (A1, mirror) and (B1, direct) reveal different spatiotemporal patterns of cortical slow waves. A2, B2. Probability of global initiation for each neuron in the network. In A1 ripples target the ‘top’ region of the cortex (cells [601-1200]) and cause higher Up-state initiation likelihood in that region (A2, blue). In B1 ripples target the ‘bottom’ region ([1-600]) and cause higher initiation in that region (B2, blue). Averages across 20 trials, dotted lines show standard error of the mean. A2 inset: Impact of ripples depends on the strength of CA1->CX connections. Color map codes probability of global Up state initiation for each neuron in a A1 wiring scenario. The preference for the upper region initiation dissolves as CA1->CX connectivity strength decreases (100% corresponds to the A2 case, 0% corresponds to cortical dynamics without CA1 input). A3,B3. The change of the Up-state initiation probability is reflected in the shape of the DUt traveling wave. A3/B3, left. Probability of DUt for each neuron as a function of time (lag) with respect to the time moment of a global DUt (zero lag). A3/B3 right. Difference of the gradient (“slope” in radians) of the DUt traveling wave in the mirror and direct map models compared to the cortex-only model (cortex receives no hippocampal input), each bar corresponds to restricted region of 100 neurons. Positive values indicate a higher tendency of waves to propagate from the bottom to the top of network when compared to the cortex-only model, while the negative values show the opposite tendency. A4,B4. Change of “incoming” synapses strength (X-axis - relative index of a presynaptic neuron in respect to the index of a fixed postsynaptic neuron) calculated using offline STDP. The neurons in the middle of the network show an opposite trend for strengthening/weakening of synapses, corresponding to preferred slope gradients as shown A3/B3. The effect starts weakening around the distance of 10 neighboring neurons (X-axis) which is due to the increasing time delay.

Preferential direction of the DUt activity propagation could have an impact on synaptic strength via spike-timing dependent plasticity (STDP). Indeed, the cortical network model used in our study reveals spindle-like activity in the beginning of each Up-state which was generated by the thalamic component of the model [Bazhenov et al., 2000, Wei et al., 2018], as is visible in the expanded timebase plot in Fig. 2 bottom left. Spindles provide a suitable time window for the STDP rule to be effective [Muller et al., 2016]. In our network model, each cortical pyramidal neuron was connected symmetrically to both sides of its close neighborhood. Thus, for each preferred direction of wave traveling, this would lead to synaptic connections decreasing strength in one direction and increasing it in the opposite direction. We estimated synaptic changes by calculating STDP offline from spike traces recorded in the simulation. Figure 6 A4/B4 shows changes to synaptic weights in close vicinity (X-axis) of each pyramidal neuron (Y-axis) for mirror (left) and direct (right) connectivity models. In the mirror model (Fig. 6A4), synapses from neurons with higher index to the neurons with lower index (corresponding to the traveling wave direction) were generally increased and opposite synapses decreased. Thus, comparing Fig. 6 A4 and B4 we found that the two wiring scenarios tend to produce symmetrically opposite synaptic changes as consequence of the opposite directions of their traveling waves.

## Discussion

The hippocampo-cortical dialogue is critical for consolidation of memory [Preston and Eichenbaum, 2013]. Coordination between the prominent oscillations during sleep – cortical slow oscillations and hippocampal ripples – was proposed to be the primary orchestrating mechanism in this dialogue [Buzsáki, 1996, Maingret et al., 2016]. In this new study, we investigated the reciprocal influence of two major sleep rhythms in the large-scale model implementing slow oscillations in the thalamo-cortical network [Bazhenov et al., 2002, Wei et al., 2018] and SWRs in the CA3-CA1 hippocampal network [Malerba et al., 2016, Malerba and Bazhenov, 2018]. Our study revealed a complex pattern of interaction between the rhythms where hippocampal ripples were able to bias cortical network to initiate Down to Up state transitions at specific cortical network locations as well as to synchronize Up to Down state transitions. This prediction may explain the mechanism behind the role of hippocampal ripples in defining cortical spike sequences replay during sleep related memory consolidation [Inostroza and Born, 2013].

Many specific details of the functional connectivity between (m)PFC and hippocampus are not fully known. Anatomically, the existence of connections from hippocampus to PFC is well established - CA1/subiculum connections to PFC were independently described in mice [Parent et al., 2009], rhesus monkey [Goldman-Rakic et al., 1984, Barbas and Blatt, 1995, Averbeck and Seo, 2008], cat [Irle and Markowitsch, 1982, Cavada et al., 1983] and in greater detail in rats (for review see Cenquizca and Swanson [2007]), where CA1 makes monosynaptic connections to (m)PFC to both excitatory and inhibitory cells [Gabbott et al., 2002] with latency in the order of 15-20 ms [Ferino et al., 1987, Laroche et al., 1990, Dégenètais et al., 2003, Tierney et al., 2004]. In line with anatomical findings, recordings during the sleep found that CA1-mPFC unit interactions were distributed widely but not uniformly across the cells and showed about 10 ms latency between CA1 and mPFC followers [Wierzynski et al., 2009]. Furthermore, there is a converging evidence that hippocampo-cortical pathway is plastic and activated during memory consolidation [Laroche et al., 2000].

Projections from PFC (anterior cingulate, ACC) back to CA1 were found in mice [Rajasethupathy et al., 2015] but parallel recording of ACC and CA1 units during SWS suggested a rather multisynaptic pathway [Wang and Ikemoto, 2016]. A weak projection between (o)PFC and CA1 was also described in rhesus monkey [Carmichael and Price, 1995] but its existence remains questionable [Cavada et al., 2000]. Thus, communication from PFC to hippocampus is likely to be mediated through the main input gate – entorhinal cortex [Lavenex and Amaral, 2000, Preston and Eichenbaum, 2013].

In agreement with these empirical studies, in our model the CA1 region projected widely into the ‘prefrontal’ cortex network, but the connectivity was not uniform, instead CA1 was parceled into small regions which targeted specific focal points in the cortical network (see diagram in Fig. 1B). In the opposite direction a small patch of the cortical network projected directly to CA3. Thus, effectively during slow-wave sleep, cortical input to the hippocampus was activated only when a traveling wave of Down to Up state transition [Massimini et al., 2004, Luczak et al., 2007, Murphy et al., 2009, Nir et al., 2011] reached that cortical patch. We need to note that because of the limited size, the model simulated global slow-wave dynamics while in vivo many Down states are isolated and may propagate only locally [Mak-McCully et al., 2015]. Without explicit modeling of dentate gyrus/rhinal cortices we considered this simplified model as a functional approximation of the intricate mechanism of how cortical input enters and affects hippocampus in vivo [Hahn et al., 2007].

Other important pathways omitted in the model included the nucleus reuniens (NR) targeting both CA1 & mPFC [Hoover and Vertes, 2012, Varela et al., 2014] and mPFC -> NR -> CA1 pathways [Vertes, 2006, Vertes et al., 2007] possibly gating mPFC->hippocampal flow which was shown to be important for memory consolidation [Ito et al., 2015].

Hippocampal SWRs were hypothesized to be a mediator of the hippocampo-cortical dialogue during deep sleep Buzsáki [1996] and indeed experimental studies revealed that cortical cells may fire in coordination with hippocampal ripples [Siapas and Wilson, 1998]. Unlike short SWR events (50-100 msec), cortical slow waves (0.2-1 Hz) are characterized by relatively smooth transitions between Up and Down states, so direct comparison of the timing of the SWR and SO events across the published studies is difficult. In many studies SWR preferably occurs during an Up-state [Mölle et al., 2006, Isomura et al., 2006, Nir et al., 2011], but see also [Battaglia et al., 2004, Hahn et al., 2007]. Another common reference points of the SO are transitions between Up and Down states; it was reported that SWRs commonly follow transition from Down to Up state in the cortical network [Sirota et al., 2003, Battaglia et al., 2004, Isomura et al., 2006, Mölle et al., 2006]. There is also evidence that SWRs precede cortical Down-state [Peyrache et al., 2009, Maingret et al., 2016, Peyrache et al., 2011].

To study interaction between SO and SWRs ripples, we used the biophysical TH-CX-HC model and observed realistic coupling behaviour between cortically generated slow oscillation and hippocampally generated ripples. In agreement with [Isomura et al., 2006], we observed clear biasing of the SWR probability by DUt; UDt transition probability increased following the ripple and causal effect of ripples was confirmed by cutting CA1->CX projections in open-loop experiment. While only a small fraction of SWRs occurred in the model during cortical Down-states, these events could still bias network location of UP-state initiation site. Open-loop (CA1->CX) simulations, where SWR was artificially triggered at different pre-defined phases of SO, revealed that the effect of SWR fundamentally depends on the phase of SO, namely we observed that i) UDt events could be both delayed or advanced by SWR; ii) synchronization of UDt events could be both improved or reduced; iii) DUt events could be only advanced; and iv) in general Up-state initiation could not be directly triggered by a single SWR event unless it occurred in very close proximity to DUt where local initiation could be observed. That is in line with experimental findings [Isomura et al., 2006] where SWR event during the Down-state was sometimes capable to trigger spiking but the network returned back to the Down state without transition to the Up-state. Similar observations led Buzsáki [2015] to suggest that in anesthesia SWR do not routinely bias the phase of slow oscillations. Our simulations suggest that while no immediate Up-state typically follows SWR events occurring during Down state, the duration of the ongoing Down state changes and thus the phase of SO may be affected as well. The model prediction that the ripple, occurring in the mid-late phase of an Up-state, is capable of advancing and synchronizing UDt, may explain data showing visible peak of ripple probability before Down-state in the UDt-triggered ripple count histogram [Peyrache et al., 2011, Maingret et al., 2016] and may shed light on the mechanisms behind apparently more synchronized UDt compared to DUt events in vivo [Volgushev et al., 2006, Chen et al., 2012]. The dependence of the SWR effect on the SO phase reported here is also in line with the experimental work of [Batterink et al., 2016], showing the existence of optimal time (with respect to SO phase) for the auditory input in targeted memory reactivation that improves memory consolidation in sleep.

We omitted in our model analysis of spindles - a very important component of the SPW-SO interaction. Studies show ripple-spindle locking [Sirota et al., 2003, Mölle et al., 2006, Clemens et al., 2007, Wierzynski et al., 2009]; a recent work reported phase-locking between SWR, SO and spindles [Latchoumane et al., 2017] and their nesting within hippocampus [Staresina et al., 2015]. While it is clear that spindle density is an important marker for memory consolidation processes [Mednick et al., 2013], the exact functional role is not clear. A recent proposal suggests SO as the leader of active memory consolidation while spindle functionally deafferenting cortical circuitry from SWR input [Genzel et al., 2014] thus helping (selective) reorganization during Up-state following SWR reactivation-Down state complex [Maingret et al., 2016].

The question about temporal coordination of DUt/UDt and SWRs is directly related to the hypothesis of the cortical and hippocampal spike sequence replay during sleep, which is believed to be necessary for stabilizing recent memory traces [Wilson and McNaughton, 1994, Nakashiba et al., 2009, van de Ven et al., 2016, Valero et al., 2017]. Cortical replay occurs during Up-state [Johnson et al., 2010], peaking close to the transition points [Isomura et al., 2006, Peyrache et al., 2011]. Hippocampal replay occurs in CA1 during the ripple events [Kudrimoti et al., 1999], it is known to be concurrent with cortical replay and both pre-cortical (PFC)[Peyrache et al., 2009] and post-cortical (visual cortex) [Ji and Wilson, 2007] coupling was observed, leading to the discussion whether the CA1 sequences are driven by SO or SWR drives replay in PFC [Genzel et al., 2014, Buzsáki, 2015]. In [Rothschild et al., 2017] it was shown that there is a bilateral dialogue between the auditory cortex (AC-CX) and CA1, where AC-CX pre-SWR firing predicted SWR content, which in turn predicted post-SWR AC-CX activity, thus suggesting a possible scenario in which a single ripple is first triggered by the cortex and intermediately influences the cortex back within a single oscillation cycle [Rothschild, 2018].

In this study, we did not attempt to model sequence replay. We observed, however, that repeated ripple events targeting a specific cortical region can reshape spatiotemporal patterns of Up-state initiation. This leads to consistent change in how Up-state waves travel across the cortical network. SWS firing patterns were found to be supportive for induction of long-term synaptic plastic changes [Chauvette et al., 2012], so we tried to assess how spike-time dependent plasticity [Bi and Poo, 2001] during slow-waves would shape the synaptic connectivity on the cortical site. We found that spatial location/distance to the cortical cells which are targets of the ripple determined whether STDP would render the synapses weakened or strengthened, thus allowing the mechanism which selects the ripple content to influence plasticity on the cortical site. Different hippocampal assemblies representing distinct sequences participating in the ripple event(s) could project into various cortical targets leading to parallel reorganization at different cortical sites. Observed reorganization by SWR is consistent with experimental results of [Maingret et al., 2016] in which proper timing of SWR (with respect to cortical Down-state) is able to reorganize mPFC firing patterns, which would otherwise stay stable [Luczak et al., 2007].

## Methods

### Hippocampal module

The model is tuned to show appropriate stochasticity in the spontaneous occurrence of sharp-wave ripples and large irregular activity in the interleaning times. The firing rates of excitatory and inhibitory cells in the system are consistent with experimental in vivo data, and the CA3-CA1 model activity in isolation from cortical input was studied in terms of the ripple mechanism [Malerba et al., 2016], the synaptic connections role on reactivation during ripples [Malerba and Bazhenov, 2018] and the relation between reactivation during sleep and synaptic changes induced by awake learning [Malerba et al., 2017b]. Technically, the model closely follows that of [Malerba et al., 2016, Malerba and Bazhenov, 2018] with a slight difference in CA3 connectivity and projections to CA1, which leads to a single main excitable region in CA3 generating all the sharp waves, rather than stochastically evolving locations in CA3 which were observed otherwise. Here we briefly describe the basic properties of the model. CA1 model consists of 800 excitatory and 160 interneurons, CA3 has 1200 excitatory cells and 240 interneurons. Each neurons is described by the following equations

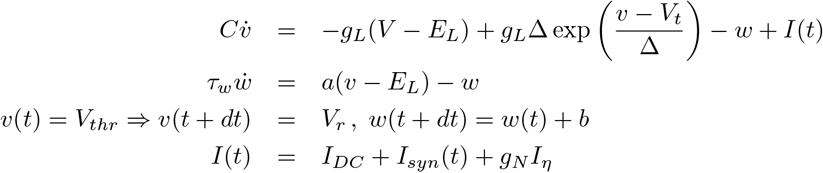

where *v* is membrane potential and *w* slow variable. For CA1 pyramidal cells *C* = 200 pF, *g_L_* = 7 nS, *E_L_* = −58 mV, Δ = 2 mV, *V_t_* = −50 mV, *τ_w_* = 120 ms, *a* = 2, *V_thr_* = 0 mV, *V_r_* = −46 mV, *b* = 100 pA; for CA3 pyramidal cells *b* = 40 pA; for both CA1/CA3 inhibitory neurons these parameter change: *g_L_* = 10 nS, *E_L_* = −70 mV, *b* =10 pA, *V_r_* = −58 mV, *τ_w_* = 30 ms.

*I_DC_* input is a constant different for each cell selected from Gaussian distribution, mean *μ* (in pA) and standard deviation *p* % expressed as a percent of the mean value was different for each population of pyramidal (*μ*_*CA*_1 = 40, *p*_*CA*1_ = 10%, *μ*_*CA*3_ = 22.5, *PcA3* = 30%) and inhibitory (*μ*_*CA*1_ = 180, *p*_*CA*1_ = 10%, *μ*_*CA*3_ = 130, *p*_*CA*3_ = 30%) cells.

We construct the background noise by generating two incoming surrogate spike trains (one excitatory and one inhibitory) and convolving each spike train with an exponential decay, and finally combining the two into a current signal *I_η_*. The two spike trains are built as memoryless, by finding the time of the next surrogate spike using an exponential random variable (note that exponential inter-arrival times are markings of memoryless Poisson Processes). To numerically obtain the exponential interarrival times of the surrogate spikes, we used a well known conversion from uniform random variables to random variables of a given distribution (inverse transform sampling). Practically, the next time a surrogate spike was generated as *R* = *t* − log(*S*)/*λ*, where *λ* = 0.5 ms^−1^ was the rate of incoming spikes, t was the current simulation time, and S a uniformly sampled random variable. Hence, we had two surrogate spike trains (with the same high rate), we convolved each of them with a exponential decay time (*τ*_*D*_ = 1 ms), shaping two noisy signals with a small standard deviation. Finally, we subtracted the inhibitory signal from the excitatory signal, and scaled the resulting signal (noisy, with one pole decay after the λ rate, and with small standard deviation) by a constant coefficient which was tuned to induce, when the current was added to a mildly hyperpolarized single cell, a standard deviation in voltage fluctuations of about 2 mV. Technically, this was obtained by scaling the noise added to excitatory cells by *g_N_* = 110.08, and to inhibitory cells by *g_N_* = 92.88.

The synaptic current from excitatory (*S_AMPA_*) and inhibitory (*S_GABA_*) synapses to neuron *n* was defined as:

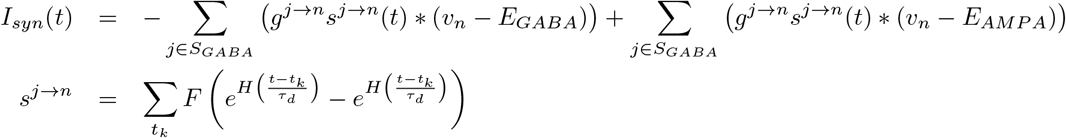

where reversal potentials were *E_GABA_* = −80 mV and *E_AMPA_* = 0 mV, *t_k_* are the spikes times from the presynaptic cell *j*. *F* is a normalization coefficient, set so that every spike in the double exponential within parentheses peaks at one, and *H* is the Heaviside function, ensuring that the effect of each presynaptic spike affects the post-synaptic current only after the spike has happened. Decay and rise constants were 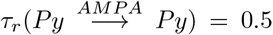, 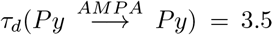, 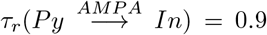, 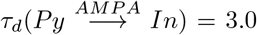, 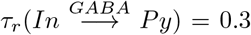, 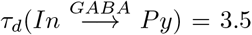, 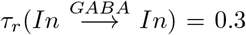, 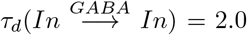; CA3 *P_y_ → In* synapses had distinct constants 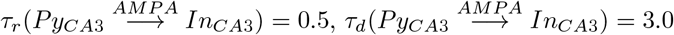.

Synaptic weights were sampled from Gaussian distributions with variance *σ* given by percent of the mean 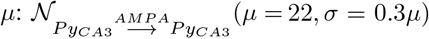, 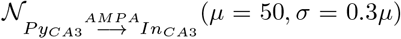, 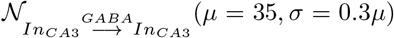, 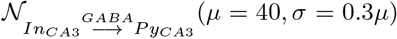, 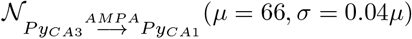, 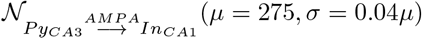.

The rationale for the choice of the equations and parameters is in length discussed in [Malerba et al., 2016, 2017a, Malerba and Bazhenov, 2018].

### Thalamo-cortical module

Our TH-CX module is similar to previous models [Bazhenov et al., 2002, Krishnan et al., 2016, Wei et al., 2018] aimed at modeling NREM stage 3 sleep. Neuromodulatory changes for different sleep stages and STDP plasticity at synapses were not employed in this model.

All neurons follow Hodgkin-Huxley kinetics, cortical neurons included dendritic and axo-somatic compartments:

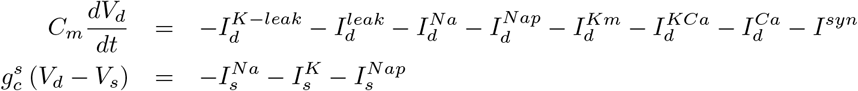

where the subscripts *s* and *d* correspond to axo-somatic and dendritic compartments. *C_m_* = 0.75 *μ*F/cm^2^, *I^K–leak^* is the potassium leak current, 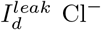 leak currents, 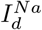 fast Na^+^ currents, 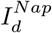 persistent sodium current, *I^K^* fast delayed rectifier K^+^ current, *I^Km^* slow voltage-dependent non-inactivating K^+^ current, *I^KCa^* slow Ca^2+^ dependent K^+^ current, *I^Ca^* high-threshold Ca^2+^ current, and *I^syn^* is the sum of all synaptic currents to the neuron. The intrinsic currents had generally the form *I^current^* = *g^current^*(*V* − *E^current^*). Details of individual currents can be found in the previous publications [Bazhenov et al., 2002, Chen et al., 2012, Wei et al., 2016]. The conductances of the leak currents were *g^K–leak^* = 0.004 mS/cm^2^ and *g^leak^* = 0.012 mS/cm^2^. The maximal conductances for the voltage and ion-gated intrinsic currents were 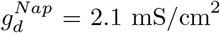, 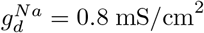, 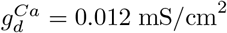, 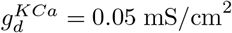, 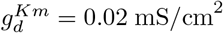, 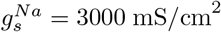, 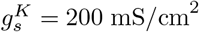, 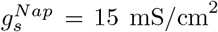. For inhibitory neurons persistent sodium current was not present (*I^Nap^* = 0) and conductances of the leak currents were: *g^K–leak^* = 0.003 mS/cm^2^ and *g^leak^* = 0.01 mS/cm^2^. The maximal conductances for the voltage and ion-gated intrinsic currents were 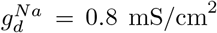, 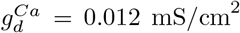, 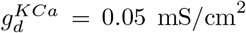, 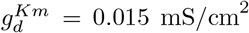, 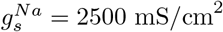, 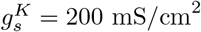.

Thalamic neurons were single-compartmental neurons following the equation:

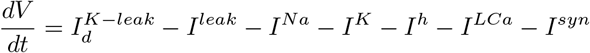

where *I^Na^* and *I^K^* are fast Na^+^/K^+^ currents, and *I^h^* hyperpolarization-activated depolarizing current. For TC neurons leak currents conductances were *g^K-leak^* = 0.035 mS/cm^2^, *g^leak^* = 0.01 mS/cm^2^, and maximal conductances for other currents were *g^Na^* = 90 mS/cm^2^, *g^K^* = 12 mS/cm^2^, *g^LCa^* = 2.5 mS/cm^2^, *g^h^* = 0.016 mS/cm^2^. For RE neurons *I^h^* current was not present (*I^h^* = 0), and conductances were *g^K-leak^* = 0.006 mS/cm^2^, *g^leak^* = 0.05 mS/cm^2^, *g^Na^* = 100 mS/cm^2^, *g^K^* = 10 mS/cm^2^, *g^LCa^* = 2.2 mS/cm^2^.

The synaptic currents for AMPA, NMDA, GABA_A_, GABA_B_ synapses were described by first order activation schemes in the form of *I_syn_* = *g_syn_*[*O*]*f*(*V*)(*V* − *E_syn_*) where *g_syn_* is maximum conductance, [O] is the fraction of open channels, *E_syn_* is the reversal potential, the details for each synaptic current is described in [Wei et al., 2016].

The maximal conductances were 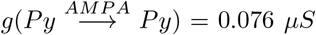, 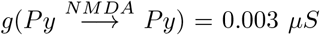, 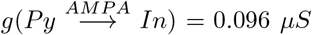, 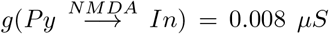, 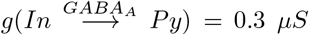, 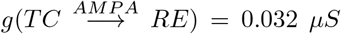, 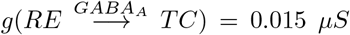, 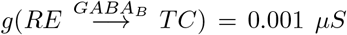, 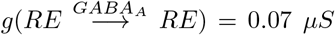, 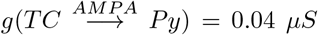, 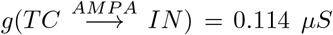, 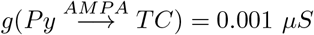, 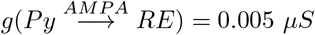.

The synapses between hippocampus and cortex were modeled as AMPA synapses from above with the possibility of signal transmission delay maximum conductances 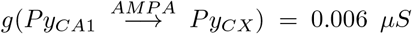, 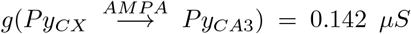, 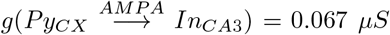, their connectivity will be described in the next section. The default synaptic delay were 16 ms [Ferino et al., 1987] for CA1->CX and 30 ms for CX->CA3 (functionally mimicking several synaptic hops needed for the cortical layer 5 signal reaching CA3 in hippocampus) connections, the small delay of CA1->CX connections did not have strong effect on the results and we set it to 0 ms later in order to have equal sampling distance when the effect of synaptic delays was explored (see Fig. 9).

Additionally in PY->PY, PY<->IN cortical connections miniature EPSP/IPSP [Redman, 1990, Salin and Prince, 1996] were present, their arrival times were modeled by Poisson processes with time-dependent mean rate *μ_AMPA_*(*t*) = (2/(1 + exp(–(*t* – *t*_0_)/*τ*)) – 1)/250 and *μ_GABA_B__*(*t*) = log((*t* – to + 50)/50)/400 with *t*_0_ is a time of last presynaptic spike, 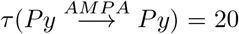, 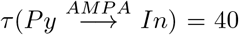. The maximal conductances for minis were 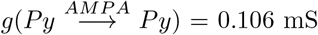, 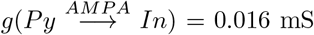, 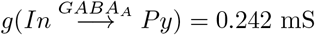.

### Network connectivity

The global topology between the different modules can be seen in Fig. 1A., the connectivity between different neuronal types can be seen in Fig. 1B. Network connectivity of thalamocortical module is similar to the previous studies [Wei et al., 2018]. Cortex (CX) consisted of 1200 layer-5 pyramidal neurons (Py) and 240 inhibitory neurons (In), thalamus consisted of 240 TC and 240 RE cells. Each neuronal type had local one-dimensional single-layer connectivity determined by the radii of connections (Fig. 1D,E). The radii of cortical and thalamic connections were 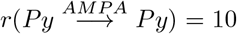, 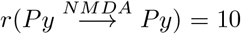, 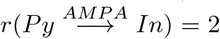, 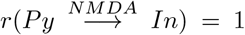, 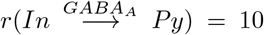, 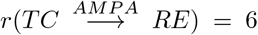, 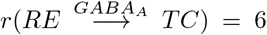, 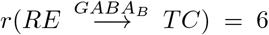, 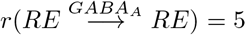, 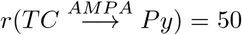, 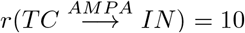, 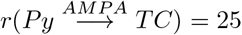, 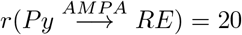. Network connectivity of hippocampal module is similar to [Malerba and Bazhenov, 2018], see Fig. 1C.

CA3 consisted of 1200 excitatory cells and 240 interneurons and had one-dimensional topology. The probability of connection from CA3 pyramidal cell with index *i* to other CA3 pyramids and interneurons was proportional to *p* = 1 – (1 – 0.15) * (*i*/1200), i.e. the subnetwork with smaller index numbers was more densely connected to increase the probability of SWR initiated in this region. The probability of connection from CA3 interneurons to CA3 pyramids was uniform, *p* = 0.7. The connectivity of Schaffer collaterals from CA3 pyramids with index *i* to CA1 pyramids and interneurons was inversely proportional, i.e. *p* = 1 – (1 – 0.15) * ((1 – *i*)/1200), so that more interconnected CA3 regions projects less to CA1 and vice versa. Within CA1 connectivity was all-to-all (including both Py and In), but synapses which were sampled at zero or less weight were removed from the connectivity, yielding to approximately 50% of CA1 Py-Py connections being cut (details in the next section).

In order to couple only the subdivision of the cortical network with hippocampus, only continuous subset of cortical pyramidal cells with index *i* ∈ [200; 399] project to subset of CA3 neurons with radii 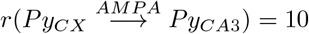, 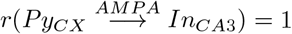. For the opposite direction flow from hippocampus to cortex, continuous chunks of 100 CA1 pyramidal neurons were targeting small continuous chunks of 5 cortical pyramids. CA1 chunks were partially overlapping (each subsequential chunk being shifted by 20 neurons), cortical chunks were non-overlapping with the gap of 33 neurons. Within the single CA1-CX chunk pair every CA1 neuron projected to all neurons of cortical chunk. The pairing between the chunks was linear and flipped, so that CA1 chunks with small index targeted chunks with high CX index and vice versa. Although by connectivity CA1 region targeted distinct loci of the whole cortical network, effectively most of the ripple input (see the histogram of firing across CA1 cells in Fig. 2, left top) was received by cortical sites opposite to the subregion projecting to CA3 thanks to the flip. Although far from real anatomical connectivity, this topology was chosen in order to be compatible with the following properties: 1. CA1 projects to frontal part of cortex (mPFC) [Cenquizca and Swanson, 2007], 2. CA1 projections target only subset of cortical cells [Laroche et al., 1990, Thierry et al., 2000, Dégenètais et al., 2003], 3. global traveling waves tend to start in the frontal region and travel towards the temporal region [Nir et al., 2011], 4. spiking activity in the temporal lobe is synaptically closer to the enthorhinal cortex, a main gate to the hippocampal structures [Ranganath and Ritchey, 2012] and is more probable to directly influence it’s activity. Translated to our connectivity, subdivision of cortical network (’temporal’ part) tends to influence SWR generation in the preferable region of CA3, which is transmitted to topologically similar region of CA1, which in turn projects to another part of the cortex (’frontal’ part).

### Computational methods

All simulations used fourth-order Runge-Kutta integration method with a time step of 0.02 ms. Custom written parallel C++ code was run on Intel Xeon Phi Cooprocessors, parts requiring exhaustive parameter search run on linux clusters through the NSG project [Sivagnanam et al., 2013]. For the basic results (Figures 3,4,6) 20 trials were run, each simulating 50 s of SWS activity. Each trial had a different random seed for initial condition of network connectivity (hippocampus) and minis generation.

At the start of the simulations, both the thalamocortical loop and hippocampal networks were allowed to run independently for a few seconds, to get their stable state before synaptically coupling the two rhythms.

Data processing was done in MATLAB (MathWorks). DUt/UDt was computed separately for each cell based on the membrane voltage, the global transition was defined as the time when most of the cells go through the transition period (peak of cells transition density). SWR event was detected for the whole CA1 network. To estimate LFP, the average synaptic current input across all pyramidal cells in CA1 was calculated, and then rescaled by 1 mS to represent a potential, such that 100pA of synaptic current produce a 100 μV LFP change. SWR was found when the filtered LFP (50–350 Hz) exceeded a threshold of 5 standard deviations of the mean computed in one SWR-free second of activity, see [Malerba et al., 2016] for details.

### Human data collection and graphoelement analysis

We obtained ~16h of NREM sleep recordings over four nights from a patient with long-standing drug-resistant partial seizures who underwent ECoG and depth electrode implantations. The depth electrode was located between the white matter and the vertical part of the hippocampal head as described in [Ding and Van Hoesen, 2015] and corresponding to image 43 of the Allen Human Brain Atlas [Ding et al., 2017].

All patients gave fully informed consent for data usage in accordance with clinical guidelines and IRB regulations at the Massachusetts General Hospital. After implantation, electrodes were located using CT and MRI [Princich et al., 2013]. Continuous recordings from SEEG depth electrodes were made over the course of clinical monitoring for spontaneous seizures, with a 512 Hz sampling rate. All SEEG contacts were converted to a bipolar transcortical montage as previously described [Mak-McCully et al., 2015]. Putative hippocampal ripples were detected following earlier published methods [Staresina et al., 2015], with some modifications to isolate sharp-wave ripples specifically: we created subject-specific average templates (−100ms to +300ms around ripple center) from hand-marked exemplars (100-400 sharp-wave ripples (SWRs) per subject) that resemble previously described primate sharp-wave ripples [Skaggs et al., 2007, Ramirez-Villegas et al., 2015]. We then evaluated whether each ripple qualifies as an SWR with the following two criteria: 1. the dot product between the template and the peri-ripple LFP (−100ms to +300ms) must be higher than those between 95% of the hand-marked SWRs and the average template. 2. the absolute difference between the LFP value at the ripple center and at the maximum/minimum value between +100ms and +250ms after ripple center was computed for each ripple; this difference must be greater than 95% of ~500 hand-marked ripples with no sharp wave. Down-states, Up-states, and slow oscillation segments in the form of down-to-up-state and up-to-down-state transitions were identified as follows: for each cortical LFP signal, a zero-phase eighth order Butterworth filter from 0.1 to 4 Hz was applied. Consecutive zero crossings within 250 to 3000 ms were then selected as putative graphoelements. For each putative graphoelement, the amplitude peak between zero crossing was computed; only the top 10% of peaks were retained. DUt and UDt transitions were then defined as centering on the midpoint between a Down/Up-state peak and the immediately following Up/Down-state peak, respectively, with the requirement that the two consecutive peaks cannot be less than 250 ms or more than 1500 ms apart.

## Acknowledgements

This work was supported by DARPA (HR0011-18-2-0021), ONR (N00014-16-1-2829-P00005), National Science Foundation (IIS-1724405).

## Supplementary info

**Figure 7:**
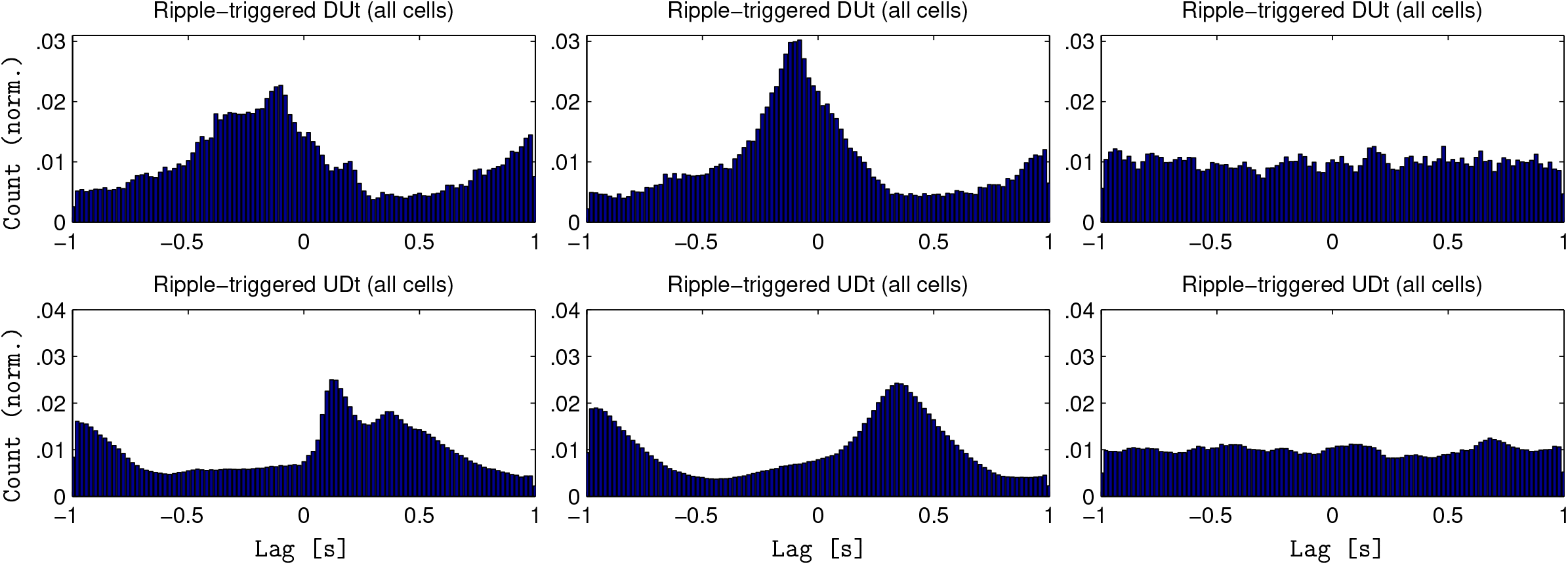
Ripple-triggered histograms. Cumulative version of heatmaps in Fig. 4., columns in the same order: left - closed loop, middle - open loop CX->HC, right - open loop HC->CX. Top. Ripple triggered DUt count (normalized). DUt of each cell is counted separately. Bottom. The same triggered by UDt.

**Figure 8:**
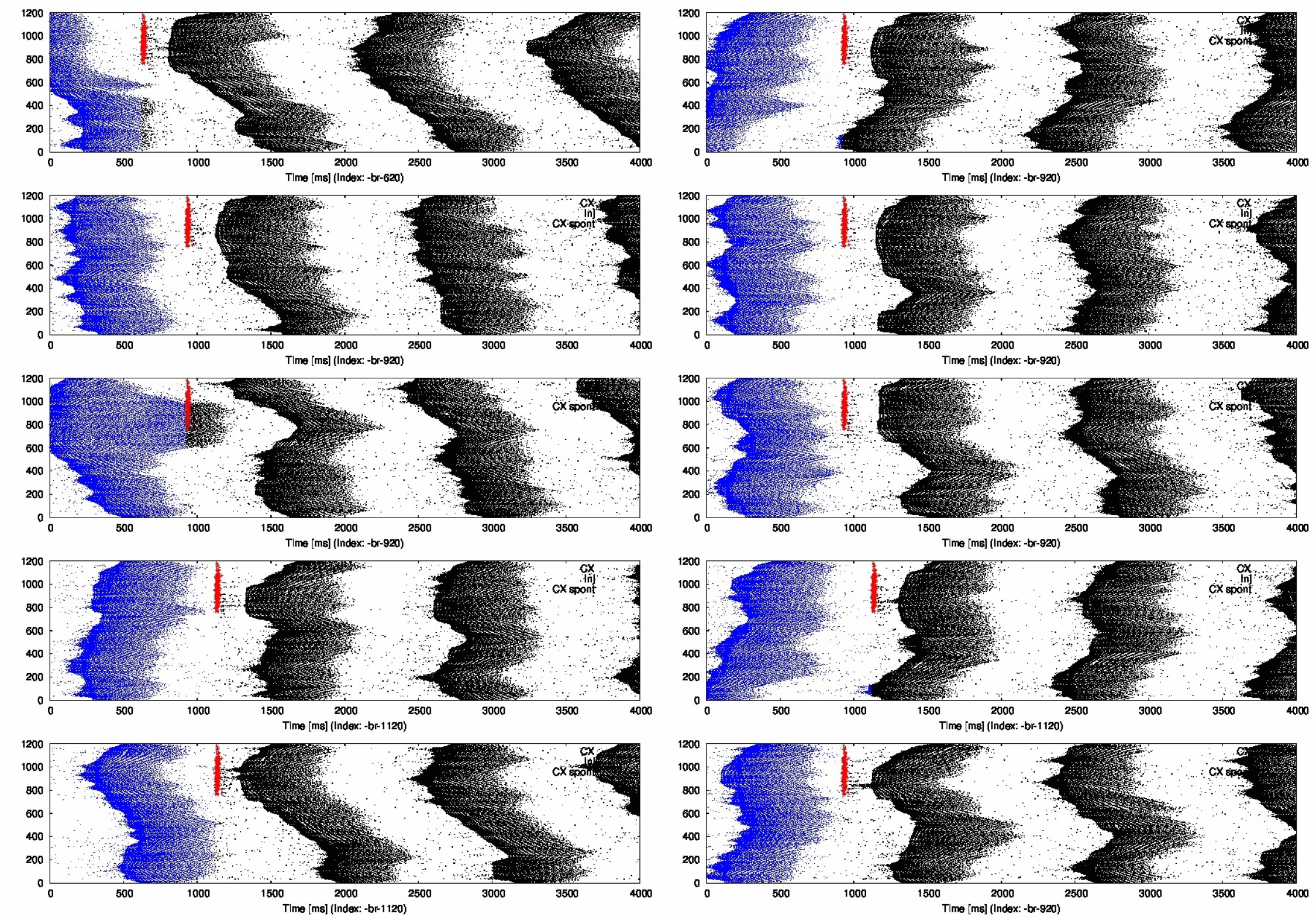
Ripple effect on the slow oscillation - animation. 10 independent runs, different random seed for each cortical run. Each run targeted by ripple (schematically interposed red) at a different phases of slow oscillation and affecting following spiking in cortex (black).

**Figure 9:**
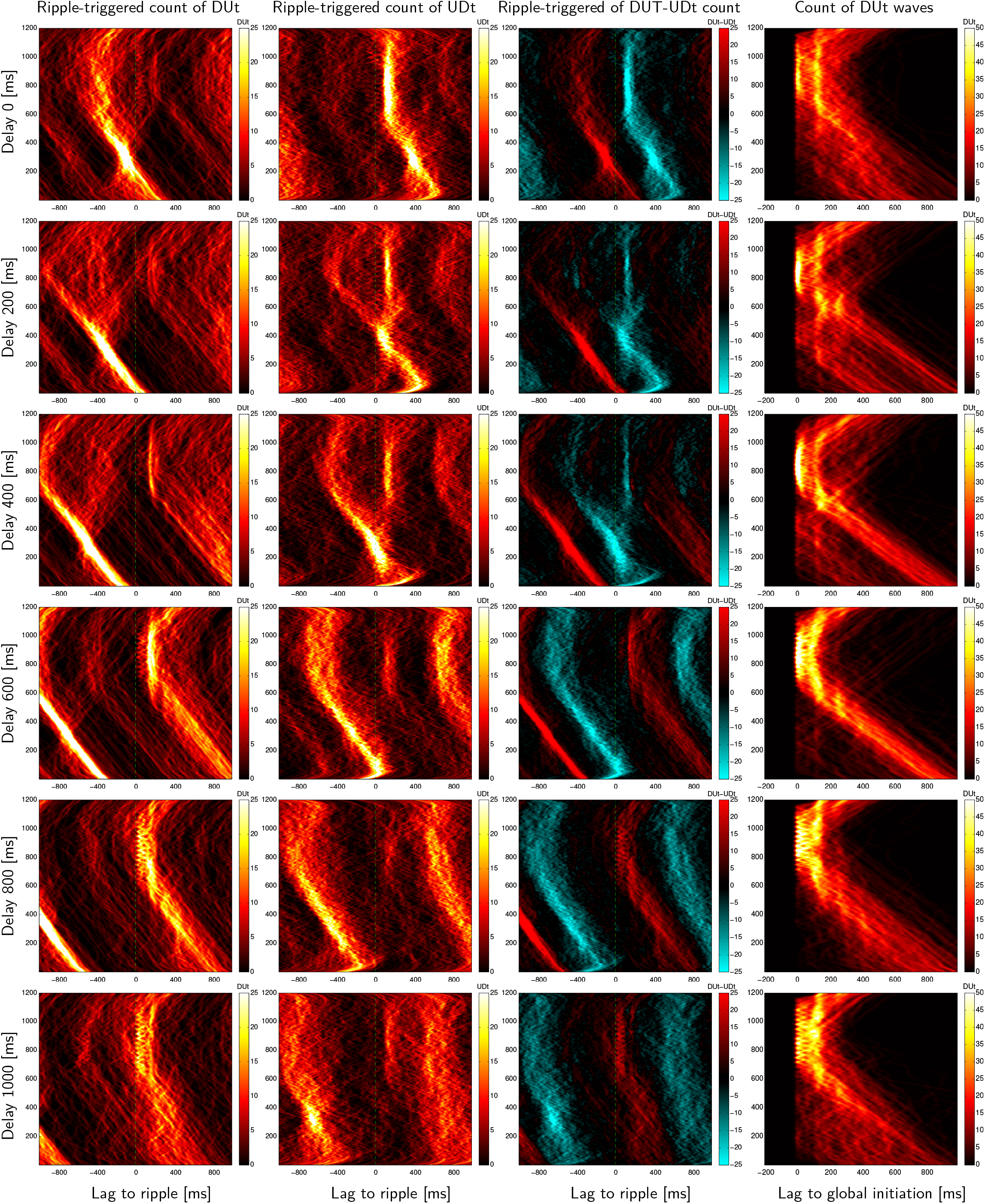
Closed-loop TH-CX-HC model run with artificially varying delay from CA1 excitatory cells to CX pyramidal cells. Individual rows show subsequent delays of 0, 200, 400, 600, 800, 1000 ms. 1st column. Spatiotemporal profile of ripple triggered DUt separately for each pyramidal cell, y-axis indexed from 1-1200 pyramidal cell, x-axis is a lag from ripple (fixed at t=0), color code shows DUt count (smoothed by gaussian kernel. Average of 10 independent trials, each 50s. 2nd column. The same for UDt. 3rd column. Aligning both DUt & UDt (DUt count – UDt count in color code). 4th column. DUt count aligned by global initiation for each DUt traveling wave.

